# BRANE Clust: Cluster-Assisted Gene Regulatory Network Inference Refinement

**DOI:** 10.1101/114769

**Authors:** Aurélie Pirayre, Camille Couprie, Laurent Duval, Jean-Christophe Pesquet

**Affiliations:** IFP Energies nouvelles, 1 et 4 avenue de Bois-Préau, 92852 Rueil-Malmaison, France.; Facebook AI Research, Paris, France.; CentraleSupélec France.

**Keywords:** Gene Regulatory Network, Clustering, Combinatorial optimization, Transcriptomic data, DREAM Challenge

## Abstract

Discovering meaningful gene interactions is crucial for the identification of novel regulatory processes in cells. Building accurately the related graphs remains challenging due to the large number of possible solutions from available data. Nonetheless, enforcing a *priori* on the graph structure, such as modularity, may reduce network indeterminacy issues. BRANE Clust (Biologically-Related A priori Network Enhancement with Clustering) refines gene regulatory network (GRN) inference thanks to cluster information. It works as a post-processing tool for inference methods (i.e. CLR, GENIE3). In BRANE Clust, the clustering is based on the inversion of a system of linear equations involving a graph-Laplacian matrix promoting a modular structure. Our approach is validated on DREAM4 and DREAM5 datasets with objective measures, showing significant comparative improvements. We provide additional insights on the discovery of novel regulatory or co-expressed links in the inferred *Escherichia coli* network evaluated using the STRING database. The comparative pertinence of clustering is discussed computationally (SIMoNe, WGCNA, X-means) and biologically (RegulonDB). BRANE Clust software is available at: http://www-syscom.univ-mlv.fr/∼pirayre/Codes-GRN-BRANE-clust.html

## 1 Introduction

**T**HE discovery of novel gene regulatory processes improves the understanding of cell phenotypic responses to external stimuli for many biological applications, such as medicine, environment or biotechnologies. To this purpose, transcriptomic data are generated and analyzed from microarrays or more recently RNAseq experiments. For each gene of a studied organism placed in different living conditions, they consist in a sequence of genetic expression levels. From these data, gene regulation mechanisms can be recovered by revealing topological links encoded in geometric graphs. Graphs and their generalizations [1] have long proved useful structures to model biological processes and to provide bioinformatics tools for data integration [2], querying [3] or Topologically Associating Domain identification [4].

### 1.1 Related works

In regulatory graphs, nodes correspond to genes. A link between two nodes is identified if a regulation relationship exists between the two corresponding genes. Such networks are called Gene Regulatory Networks (GRNs). Their construction as well as their analysis remain challenging despite the large number of available inference methods [5], [6], [7], [8]. Metric- or model-based are the principal approaches to infer a GRN. This first one (metric-based) roots on the computation of statistical measures reflecting the similarity or the dependence between pairwise gene expression profiles. Mutual information is the most common metric: Relevance Network [9], CLR [10], MINET [11] or ARACNE [12]. The second one relies on different models. Differential equations are used on *time-course* data [13]. Graphical models (GM), such as (probabilistic) Boolean or Bayesian networks, are particularly well investigated [14], [15], [16], [17], [18]. For instance, they estimate the covariance matrix, directly providing the GRN from its inverse [19]. Integrating additional knowledge on miRNAs, authors in [20] determine both transcriptional and post-transcriptional regulations through dynamic Bayesian network. In the recent work of [21], authors developed the RegNetwork tool to provide GRNs by combining knwoledge from various databases such as TRED, KEGG or TFBS, for instance. GENIE3 uses random forests to solve a set of regression problems [22]. Methods that provide a complete weighted network require a selection of meaningful relationships between genes (GRN), ideally relevant to biological interpretation. This is classically done by removing edges whose weights are lower than an arbitrary value. However, due to the very limited length of expression signals with respect to the number of genes, satisfactory results are difficult to obtain. Incorporating *a priori* in GRN estimation leads to more reliable and interpretable results. For instance, [23] uses both node and edge information: genes coding for transcription factors (TFs) and co-regulatory relationships. GRACE [24] models *a priori* and modules using multiple heterogeneous data integration with random forest regression and Markov Random Fields.

Network construction is often decoupled from further analysis, including module detection as in WGCNA [25]. However, one can benefit from the incorporation of modular structures at earlier stages of GRN inference. Thus, compounding inference and clustering more directly can better take network topology into account [26]. To the best of our knowledge, only very few methods perform joint clustering and graph inference. They often rely on graphical models to improve the network inference task. While authors in [18] use Gaussian Graphical Models (GGM) with a penalization integrating a latent structure, [14] uses cluster results as a GGM preprocessing. An iterative module learning procedure, based on the Expectation-Maximization algorithm, is proposed by [27] to deal with a probabilistic graphical model. This procedure is improved in [28] using an ensemble of possible statistical models. Authors in [29] infer regulatory programs probabilistically constrained to reveal a modular organization of the underlying regulatory network.

Despite combining inference and clustering in interesting frameworks, the scarce ratio between observations (experimental conditions and replicates) and variables of interest (genes) in real data generally enfeebles their probabilistic model usage.

### 1.2 Our contributions

Contributions are threefold, on GRN refinement formulation, resolution and validation.

#### Formulation

BRANE Clust embeds *soft*-clustering toward network inference refinement. Allowing cluster fusion, it generalizes [30], limited to one cluster per TF. A combinatorial optimization approach primarily generates a topology from a complete weighted graph, and segments genes into clusters. The chosen energy to be minimized consitutes a significant refinement of the one used in [23].

#### Resolution

A fast alternating optimization method achieves joint inference and clustering. The optimal binary variables on the edges define the graph structure. They are obtained *via* an explicit solution. Optimal clustering defined by variables on nodes is computed by the *random walker* algorithm.

#### Validation

Extensive experiments are performed on eight networks of diverse types and sizes (DREAM4 and DREAM5). Inference and clustering pertinence are discussed. Computational comparisons with CLR, GENIE3, SIMoNe, WGCNA, X-means are led, and parameter sensitivity is analyzed. Biological interpretation on the *Escherichia coli* (*E. coli*) network and the estimated gene clusters is carried out respectively with STRING and RegulonDB databases.

Section 2 expresses classical edge selection as an optimization problem and its cluster-assisted refinement. *Hard*-clustering [30] is recalled and its generalization (*soft*-clustering) is then detailed as well as the proposed computational strategy. BRANE Clust (*hard* and *soft*) is evaluated and discussed in Section 3, followed by a conclusion in Section 4.

## 2 Problem formulation

### 2.1 GRN extraction as edge selection optimization

A graph structure may represent gene regulation from expression data. Let 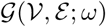 be a fully connected undirected, weighted graph. The set of gene indices is 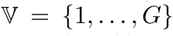. The set of nodes (corresponding to genes) is 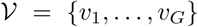. The set of edges ***ε*** has cardinality *n* = *G*(*G –* 1)/2, and *ω* denotes edge weights

Weight *ω_i_*,*_j_* is assigned to each edge *e_i,j_*,(*i,j*) 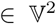. It reflects the level of interaction between genes *i* and *j*. Microarray or RNA-seq experiments produce gene expression profiles. They yield graph weights, using for instance CLR or GENIE3. A GRN is classically extracted from 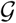 by removing edges whose weights are lower than a threshold λ. Elaborating on [23], this graph thresholding step is formalized with a binary labeling *x_i,j_* of edges *e_i,j_* as:

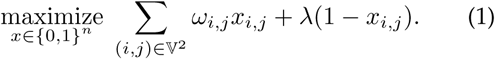

In (1), the first term alone would select all edges. The second term restricts this selection to those with weights larger than λ. The optimal labeling is given by the explicit solution:

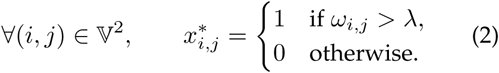

Binary labels define edge presence indicators by setting *x_i,j_* to 1 if the edge *e_i,j_* is in the final graph 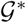 and 0 otherwise.

Lowering λ increases the potential of recovering known gene interactions. However, unassisted threshold selection may unveil an excessive number of false positives in the GRN. We thus integrate a clustering step in the graph thresholding formulation to refine the GRN toward an increased reliability. We now unfold and generalize the cluster-assisted approach proposed in [30].

### 2.2 Cluster-assisted graph inference

Relying on sound gene clustering, one can better control a modular graph structure. We consider TF-centric modules as groups of genes arranged around transcription factors. This additional knowledge is used for prediction, as TF-centric modules favor the detection of new target genes.

We thus integrate a clustering step into (1). It promotes the presence of edges linking nodes belonging to the same cluster. We want to design a cost function so as to impact weights in (1) as follows. If nodes *υ_i_* and *υ_j_* belong to:

- the same cluster, weights remain unchanged,
- distinct clusters, weights are reduced.

Let 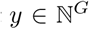 denote a node cluster labeling vector. Let 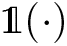 denote the characteristic function equal to 1 if its argument is verified and 0 otherwise. A parameter *β* > 1 is used to control the clustering. An instance of cost function is:

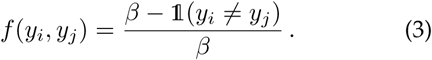

If nodes belong to the same cluster, *f*(*y_i_, y_j_*) = 1 independently of *β*. If nodes belong to different clusters, *f*(*y_i_*,*y_j_*) may vary from 0 for low *β* to 1 for high *β*, thus emulating standard thresholding. Figure 1 illustrates the effect of the clustering in the inference process.

**Figure 1.**
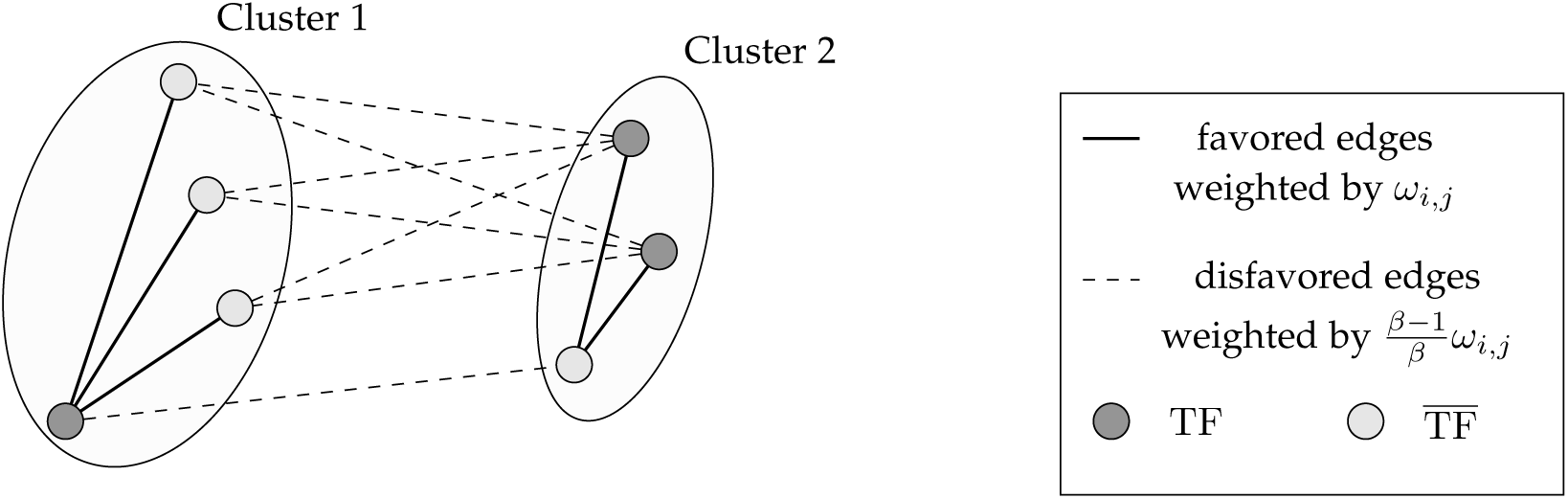
Schematic view of the clustering integration in the inference.

We assume given *T* central nodes, for instance gene hubs [26]. We choose them here as the list of genes known to code for TFs, 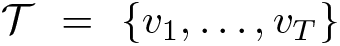, indexed by the set 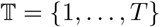.

We now present two cluster-assisted approaches to improve network inference, based on *hard*- and *soft*- clustering, the second encompassing the first.

### 2.2.1 Hard-clustering

We firstly propose a modular structure tightened by a unique and distinct TF per cluster. We elaborate the edge problem (1) into a node-and-edge optimization:

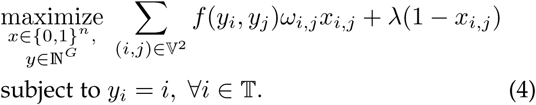

In this case, the GRN inference is assisted by *hard-*clustering: the constraint on the labeling *y* is enforced exactly. Indeed, the proposed constraint is equivalent to pre-labeling TFs and to assigning labels to non transcription factors 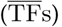. In the graph, the linkage of 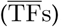 (belonging to cluster *i*) to the corresponding TF *υ_i_* is fostered. Here, the number of clusters is set to *T* in an *ad hoc* manner and admits a closed form solution [30].

### 2.2.2 Soft-clustering

While (4) yields good results on simulated data, it can be softened to better mimic biological scenarios. A possible generalization is

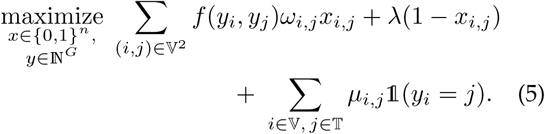

Set *μ_i_*,*_j_* = α when *i* = *j* and zero elsewhere. Letting α → ∞ recovers the *hard*-clustering solution(4).

With α finite, complementing *μ_i_*,*_j_*· weights for *i* ≠ *j* promote the merging of TF-centric clusters. Fusion is driven by strong-enough relations between TF *j* and any gene *i*. A level *τ* ∈ [0, 1] conditions the merging criterion defined by 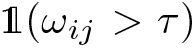. It is weighted differentially with the nature of gene *i*. When 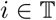 a large α factor promotes TF-centric modules. When 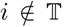, the 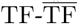 interaction is simply weighted by ω*_ij_*, to preserve an influence of potentially undiscovered TFs. Consequently, we set:

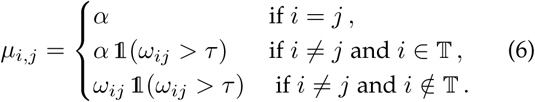

Intuitively, cluster merging depends on strong-enough TF-gene relations. We subsequently fix α to their cardinality: 
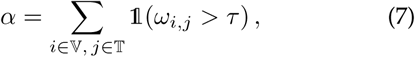
which is consistent with the order of magnitude optimal parameters obtained experimentally.

We now describe the optimization strategy for *soft*-cluster-assisted graph inference.

### 2.3 Optimization strategy

Problem (5) can be split into two sub-problems. *Soft-*cluster-assisted inference is then solved through an alternating optimization scheme. At fixed *y* and variable *x*, Problem (5) becomes:

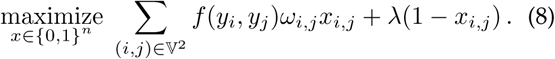

Its solution is explicit:

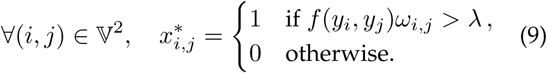

At fixed *x* and variable *y*, Problem (5) reduces to 
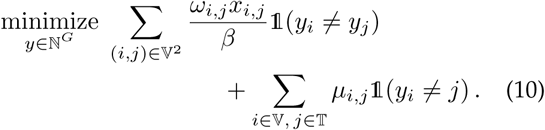

Independently from the choice of *μ_i,j_* i.e. *hard-* or *soft*-clustering, the optimization of Problem (10) is illustrated with the graph structure of Figure 2. As displayed by Figure 2(b), the presence of strongly weighted edges between two TFs favors their merging. Merging is also possible for 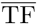 genes that also exhibit a strong weight with a TF. This copes with the fact that not all TFs are known in real biological datasets.

**Figure 2.**
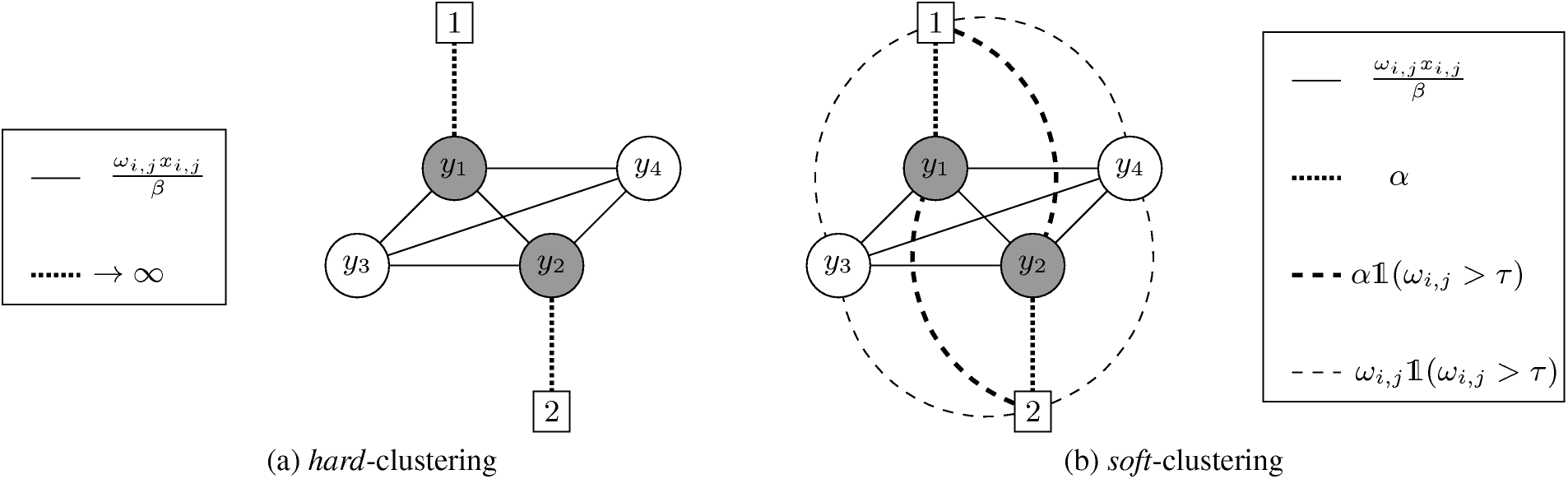
Graph built to perform (a) *hard-* and (b) *soft*-clustering. We define as many markers as TFs (gray nodes), labeled with TF indices. In the hard-clustering case, optimization constrains all node labels *y* to be equal to one of the markers. For *soft*-clustering, thanks to weights *μ_i,j_*, two clusters are merged if their respective TFs have strong weights. Legend-box *α* parameter for *soft*-clustering refers to (7).

Unfortunately, the cost function in (10) is NP-hard. It can be harnessed with random walker algorithm [31]. Cluster labels are obtained by relaxing simpler binary sub-problems. Binary label values relaxed in [0, 1] are interpreted as probabilities. Maximally probable outcomes finally yield optimal cluster labeling.

We seek clusters attached to TFs. This label restriction to 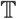 is tackled by defining the set {*s*^(1)^,…, *s*^(*T*)^}, with *T* binary vectors of length *T*. To emulate the second term in (10), their components are set to 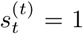 and 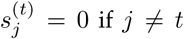. Let 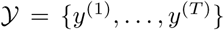 be a set of *T* vectors. For all 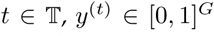 contains the probabilities for nodes to be assigned to cluster *t*. Problem (10) is re-expressed as: 
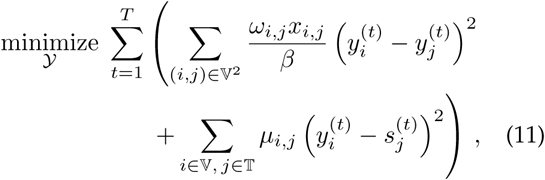
which can be decomposed into *T* sub-problems. A given sub-problem *t* evaluates *y*^(^*^t^*^)^ with respect to vector *s*^(^*^t^*^)^. Its graph interpretation resorts to fixing marker label *t* to 1 and the others to 0, as described in Figure 3(b). Probability 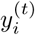 reflects the chance to reach the marker labeled by 1 first, for a random walker leaving node *i* in the graph. Higher weights encode preferable paths for the walker, and therefore drive the computed probabilities.

**Figure 3.**
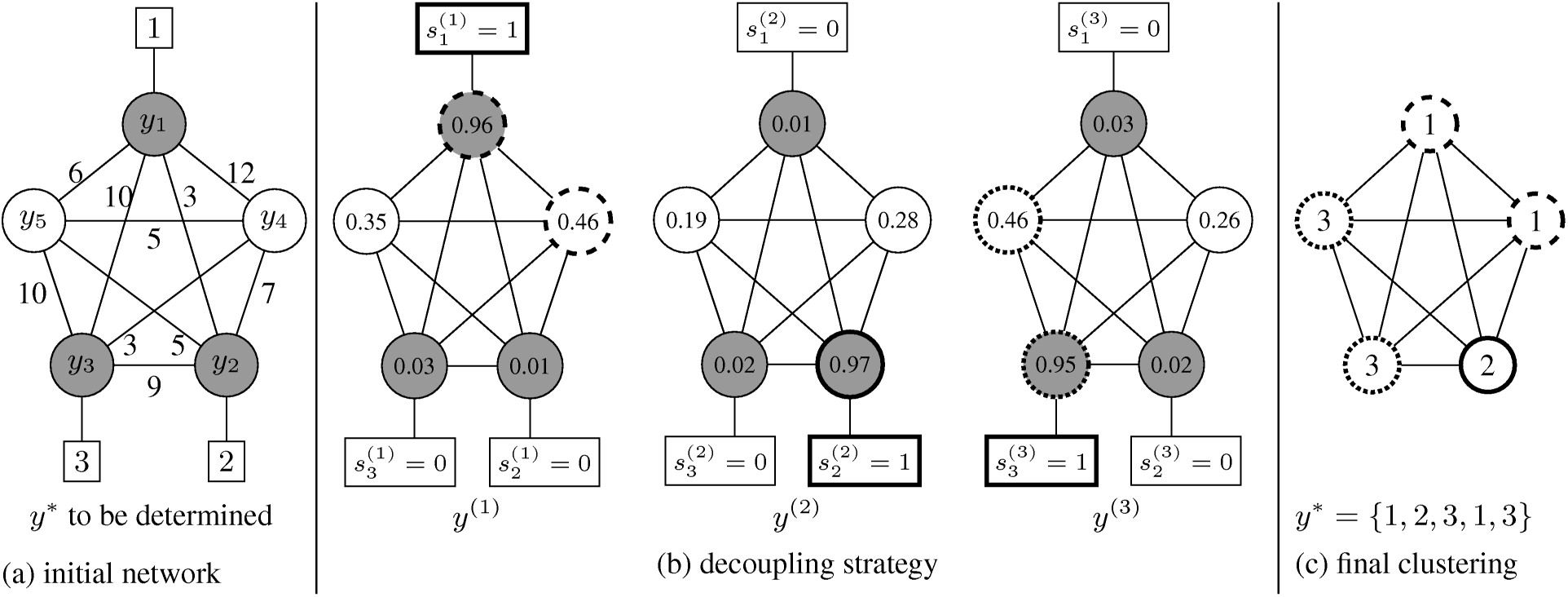
Graph-based clustering using a decoupling strategy for *hard*-clustering. The principle is similar for *soft*-clustering. Gray nodes represent TFs nodes. The *T*-label problem is decomposed into *T* binary sub-problems by setting the component *t* of marker labels *s*^(*t*)^, 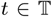, to one and the others to zero. Each sub-problem *t* leads to a probability for each node. The final node clustering corresponds to the label whose probability amidst the *T* sub-problems is maximal.

Formulation (11) is an instance of the combinatorial Dirichlet problem. The energy of the latter involves the Laplacian matrix defined as:

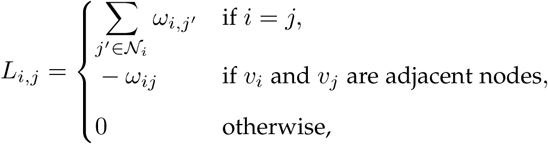
Where *𝒩_i_* denotes the set of indices of the nodes adjacent to *v_i_*. The energy writes, in its simplest form, 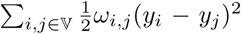, which can also be expressed as 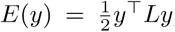. Equation (11) follows the same pattern with the Laplacian matrix of the graph in Figure 2(b). The minimization of (11) amounts to solving *T* – 1 systems of linear equations admitting a unique solution. The maximum probability arising from sub-problem 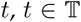, defines each node label as illustrated in Figure 3(c). The optimal cluster labeling 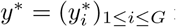 is given by

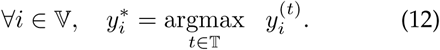

An approximate solution to Problem (5) yields the GRN after few iterations of alternating optimization between (8) and (11), less than 20 with our datasets.

## 3 RESULTS AND DISCUSSIONS

We first explain the evaluation and comparison methodologies. We then provide extensive numerical results, present a thorough interpretation on *E. coli* and discuss issues on inference and clustering.

### 3.1 overall evaluation methodology

#### 3.1.1 Metrics and methods

BRANE Clust is evaluated by comparing inferred networks to reference ones on several *in-silico* and real microarray datasets from DREAM4 [5] and DREAM5 [6] challenges. Evaluations are based on the computation of Precision (P) and Recall (R) measures. Their computation can be performed in different fashions. DREAM4 or DREAM5 estimate them based on weight ranking on complete graphs. This cannot be used in our context since BRANE Clust infers a binary-valued network. We therefore opted for the method proposed in [10]. If #TP, #FP and #FN respectively denote the number of True Positive edges (in both inferred and reference networks), False Positive (inferred but not in reference network) and False Negative (not inferred but in reference network), P and R are defined by:

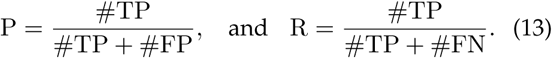

GRN construction methods are then compared by constructing Precision-Recall (PR) curves. The latter are obtained by spanning the same range of thresholds for all the compared methods. Since we use normalized weights, λ is chosen to vary linearly between 0 and 1. Several methods — classical thresholding, BRANE Cut, BRANE Clust (*hard* and *soft*)— are finally ranked according to the highest Area Under the PR curve (AUPR). We compare them on two kinds of input weights obtained from state-of-the-art inference methods: CLR [10] and GENIE3 [22]. Both methods integrate knowledge about TFs. These two methods are frequently used as benchmarks [11], [13], [23], [29]. Notably, GENIE3 was the best performer in the used DREAM4 and DREAM5 datasets.

#### 3.1.2 Parameter settings

DREAM4 *in-silico* data is generated with GeneNetWeaver [32], [33] and is based on true networks. A perfect knowledge of TFs is thus available and simulated gene expressions are considered more reliable. Hence, we have more confidence in strong edge weights for the cluster fusion task. Higher merging criterion levels *τ* are therefore preferred to drive cluster fusion on simulated data. With real data, conversely, uncertainty in experimental gene expressions and partial knowledge of TFs are an incentive for lower levels. The latter ones tend to redeem lower weights, affected by experimental biases and variability. Therefore, *τ* is set to either 0.8 (*in-silico* datasets) or 0.3 (real datasets). Satisfactory trade-offs are obtained with the same parameter setting for all experiments, whatever the data (size, weights, number of TFs) and the methods. Clustering influence on the inference is controlled by *β* = 2. The impact of *τ* and *β* variations on inference is assessed by a sensitivity analysis.

In practice, the moderate value of *β* = 2 is an efficient initial choice. Finally, a suitable range for *τ* resides around the central inter-quartile range. In other words, *τ* values close to 0 or 1 would disfavor either clustering or inference, unbalancing the performance of BRANE Clust. Globally from novel data, the order of magnitude of *τ* could be determined first by setting *β* = 2, then marginal improvements can be obtained by adjustment from this starting point.

### 3.2 Numerical results

#### 3.2.1 Results on DREAM4

The DREAM4 multifactorial challenge [5] is composed of five datasets, corresponding to simulated gene expression data. Each dataset is composed of 100 genes simulated in 100 conditions. The validation procedure uses the reference graph for each network. We first compute the classical thresholding reference on CLR and GENIE3 weights (T-CLR and T-GENIE3). We then refine these results with BRANE Cut and BRANE Clust (*hard* and *soft*). Table 1 gathers corresponding AUPR and gains. It demonstrates the advantage of BRANE Clust over both classical thresholding and BRANE Cut, and of the *soft* over the *hard* version.

**Table 1.** DREAM4 *in-silico* dataset: AUPR for T-CLR, T-GENIE3, BRANE Cut, BRANE Clust (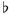: *hard* or ^#^: *soft*-clustering). Percentage gain are over classical thresholding.

| Network | 1 | 2 | 3 | 4 | 5 | Average |
| --- | --- | --- | --- | --- | --- | --- |
| T-CLR | 0.256 | 0.275 | 0.314 | 0.313 | 0.313 | 0.294 |
| BRANE Cut | 0.282 | 0.308 | 0.343 | 0.344 | 0.356 | 0.327 |
| Gain | 10.1 % | 11.8 % | 9.1 % | 9.9 % | 13.7 % | 10.9 % |
| BRANE Clust<sup>b</sup> | 0.291 | 0.288 | 0.358 | 0.356 | 0.355 | 0.330 |
| Gain | 13.7 % | 4.7 % | 14.0 % | 13.7 % | 13.4 % | **11.9 %** |
| BRANE Clust<sup>#</sup> | 0.275 | 0.337 | 0.360 | 0.335 | 0.342 | 0.330 |
| Gain | 7.3 % | 22.6 % | 14.5 % | 7.0 % | 9.1 % | **12.1 %** |
| T-GENIE3 | 0.269 | 0.288 | 0.331 | 0.323 | 0.329 | 0.308 |
| BRANE Cut | 0.298 | 0.316 | 0.357 | 0.344 | 0.352 | 0.333 |
| Gain | 10.7 % | 9.9 % | 7.8 % | 6.5 % | 7.0 % | 8.4 % |
| BRANE Clust<sup>b</sup> | 0.286 | 0.313 | 0.386 | 0.360 | 0.369 | 0.343 |
| Gain | 6.3 % | 8.7 % | 14.2 % | 11.4 % | 12.1 % | **10.5 %** |
| BRANE Clust<sup>#</sup> | 0.287 | 0.348 | 0.364 | 0.371 | 0.367 | 0.347 |
| Gain | 6.5 % | 20.9 % | 10.0 % | 15.0 % | 11.6 % | **12.8 %** |

Initialized either with CLR or GENIE3 weights, BRANE Clust (*soft*) ameliorates simple threshold results (Table 1). An improvement is observed over the five networks: about 12% using CLR and 13% with GENIE3. Best results are obtained on Network 2, with gains of about 23 % (CLR) and 21 % (GENIE3). Improvement is higher for networks with low Recall and high Precision. They are often composed of a relatively low number of edges (less than 250). Comparisons with BRANE Cut demonstrate the advantage of the clustered gene groups *a priori* over a mere co-regulation. BRANE Clust (*soft*) gains are greater in percentage points by 1.2 p.p. and 4.4 p.p. on CLR and GENIE3, respectively. In addition, *soft* version provides more favorable results than *hard* version of BRANE Clust with average gains of 0.2 p.p. and 2.3p.p. on CLR and GENIE3, respectively. In the general case, BRANE Clust (*soft*) provides networks with better predictability, leading to a more reliable interpretation.

#### 3.2.2 Sensitivity analysis

BRANE Clust *soft* parameters *β* and *τ* are fixed in all our validations according to Section 3.1.2. However, they could affect inference results. A sensitivity analysis, based on a grid-search strategy, evaluates their impact. The parameter *τ* controlling cluster fusion varies between 0.1 and 0.9 with a 0.1 step. The *β* parameter controlling clustering influence in the inference varies between 1.1 and 2 with a 0.1 step, and between 2 and 5 with a unit step. AUPR for each couple of parameters are computed and results are compiled in Figure 4. For each *τ*, we report the average AUPR and its standard deviation over *β*. Globally, on the five networks and with both CLR and GENIE3 weights, we observe that, except for only few cases, the average AUPR is significantly higher than the reference (T-CLR and T-GENIE3, Section 2.1). Although the variability over *β* often increases with *τ*, higher *τ* yield significantly better AUPR. The increase in *β* variability with *τ* may be explained by the selectivity of cluster merging. Low *τ* levels significantly trigger cluster fusion. The reduction in the number of labels diminishes the impact of *β*

**Figure 4.**
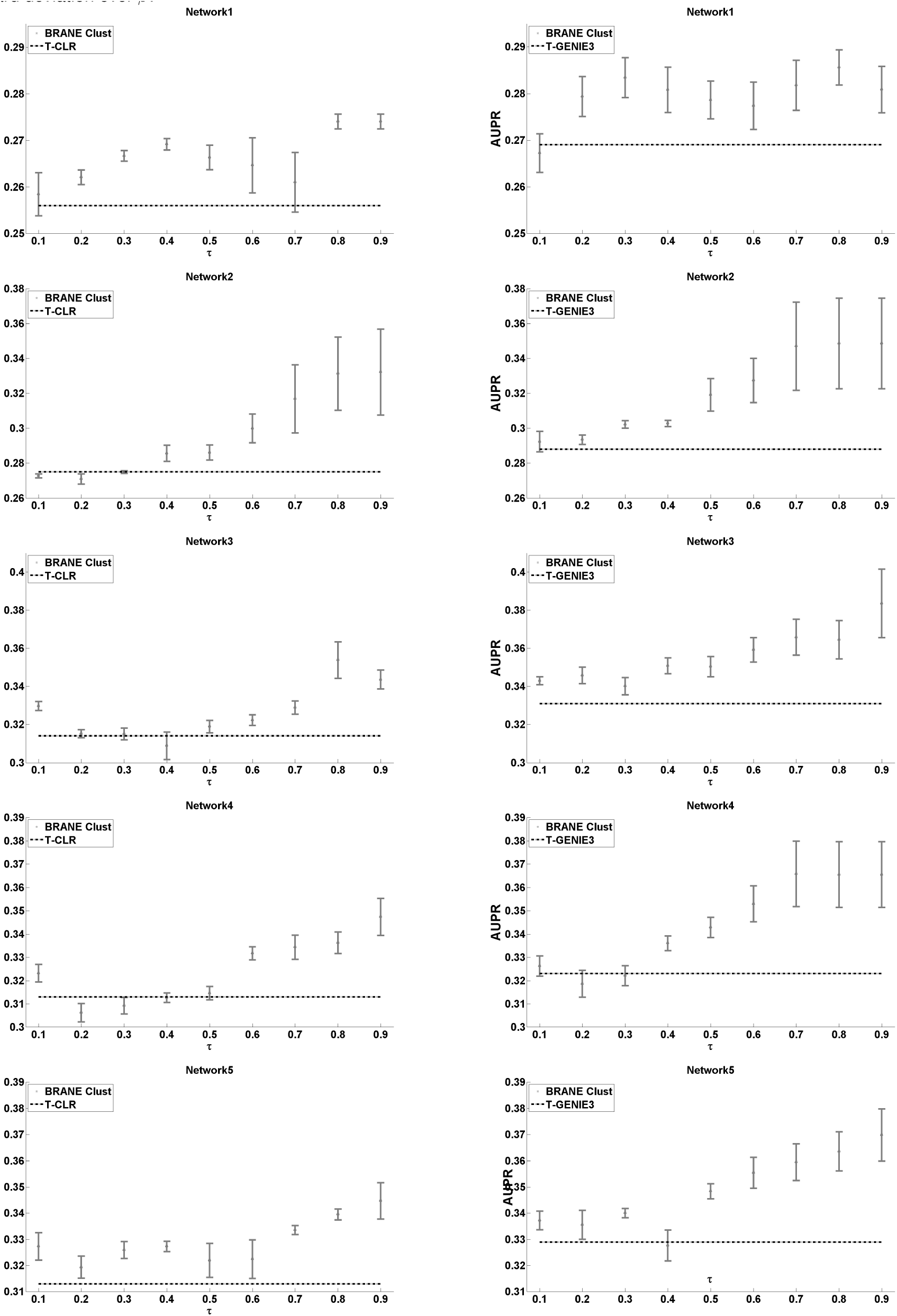
Grid-search-based sensitivity analysis of parameters *β* and *τ*. For each *τ*, results are given in terms of average AUPR and standard deviation over *β*.

#### 3.2.3 Results on DREAM5

Gene/condition and TF/gene ratios for DREAM4 networks do not reflect real datasets properties. A second evaluation is therefore performed on DREAM5, containing more realistic data. This challenge [6] is composed of four networks. Three networks only (1, 3 and 4) present an associated ground truth. The synthetic dataset 1 is composed of 1643 genes (195 TFs) and 805 conditions. The two others (3 and 4) come from real data: *Escherichia coli* and *Saccharomyces cerevisiae*, respectively. The *E. coli* dataset contains 805 conditions for 4511 genes (334 TFs). The S*. cerevisiae* dataset encompasses 5950 genes (333 TFs) and 536 conditions. AUPR for these networks are given in Table 2 with the corresponding curves in Figure 5.

**Table 2.** DREAM5: AUPR for T-CLR, T-GENIE3, BRANE Cut, BRANE Clust (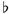:*hard* or ^#^:*soft-* clustering). Percentage gains are over classical thresholding.

| Network | 1 | 3 | 4 |
| --- | --- | --- | --- |
| T-CLR | 0.252 | 0.0378 | 0.0080 |
| BRANE Cut | 0.269 | 0.0388 | 0.0079 |
| Gain | 6.7 % | 2.8 % | - |
| BRANE Clust<sup>b</sup> | 0.309 | 0.0270 | 0.0077 |
| Gain | 22.4 % | -28.5 % | - |
| BRANE Clust<sup>#</sup> | 0.253 | 0.0399 | 0.0073 |
| Gain | 0.4 % | 5.5 % | - |
| T-GENIE3 | 0.283 | 0.0488 | 0.0081 |
| BRANE Cut | 0.297 | 0.0498 | 0.0083 |
| Gain | 4.9 % | 2.1 % | - |
| BRANE Clust<sup>b</sup> | 0.328 | 0.0383 | 0.0083 |
| Gain | 16.0 % | -21.5 % | - |
| BRANE Clust<sup>#</sup> | 0.327 | 0.0536 | 0.0083 |
| Gain | 15.5 % | 9.8 % | - |

**Figure 5.**
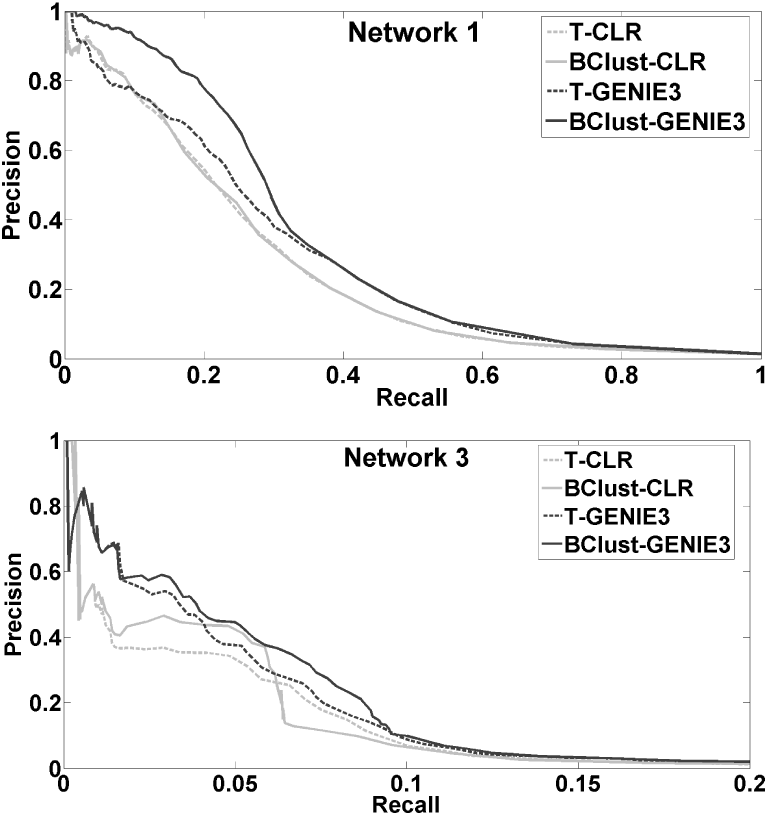
DREAM5: PR curves for T-CLR, T-GENIE3, and BRANE Clust^#^, on networks 1 (top) and 3 (bottom).

**Figure 6.**
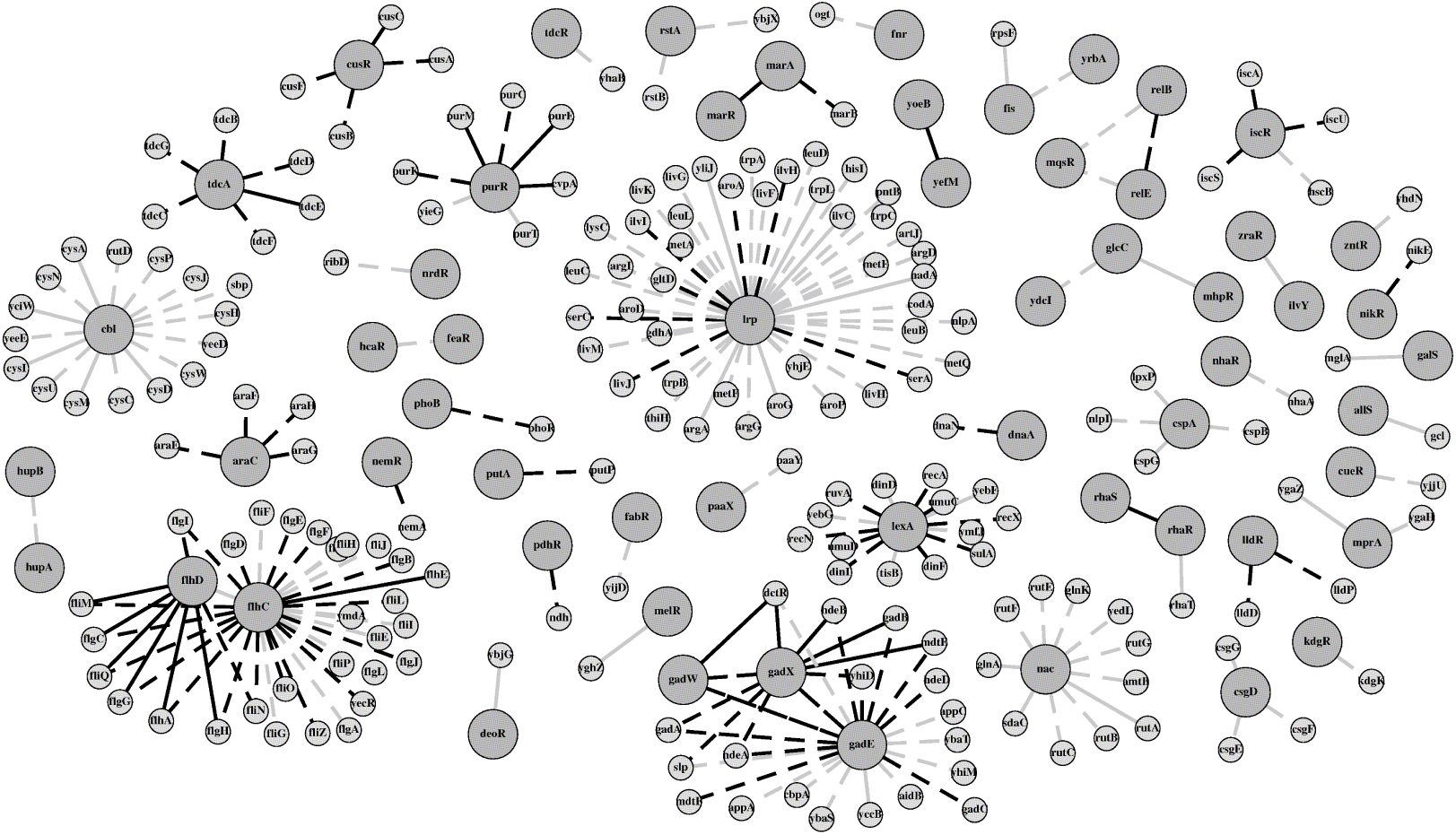
Network built using BRANE Clust on GENIE3 weights and containing 236 edges. Large dark gray nodes refers to transcription factors (TFs). Inferred edges also reported in the ground truth are colored in black while predictive edges are light gray. Dashed edges correspond to a link inferred by both BRANE Clust and GENIE3 while solid links refer to edges specifically inferred by BRANE Clust.

The tininess of the AUPR for both CLR and GENIE3 on Network 4 impedes a relevant interpretation of gains using BRANE Clust. Results on Network 1 show small refinement on CLR but a 15% improvement on GENIE3 weights. Improvements on real *Escherichia coli* experiments (Network 3) are higher for CLR, with a gain of about 6 %. The gain using GENIE3 weights is close to 10 %. The best improvements dwell within different parts of the PR curves: high precisions for simulated data, intermediate precisions for real data.

Table 2 also presents comparisons with previous network thresholding post-processing, BRANE Cut and BRANE Clust (*hard*). Results on Network 1 show the highest improvement with BRANE Clust (*hard*) on CLR and with BRANE Clust (*hard* and *soft*) on GENIE3. Moreover, on real data (*E. coli*), we found that BRANE Clust (*soft*) offers better results. On Network 3, comparisons with BRANE Cut show that BRANE Clust (*soft*) gains are greater by 2.7 p.p. and 7.7 p.p. on CLR and GENIE3, respectively. As expected, the limitation to one TF per cluster in BRANE Clust (*hard*) may be harmful on real data. BRANE Clust (*soft*) provides gains greater by 34 p.p. (CLR) and 31.3 p.p. (GENIE3) compared to BRANE Clust (*hard*). While the interest of gene cluster merging may be discussed on simulated data, its advantage on real data is demonstrated by significant gains on objective measures for the *E. coli* network.

Nevertheless, as shown in Figure 5, performance of BRANE Clust can become quite competitive with respect to classical thresholding. This observation specifically makes sense for extreme λ values. Indeed, when λ is high (low Recall and high Precision), the final network is critically sparse, yielding to a negligible influence of the clustering in the inferred network. Conversely, when λ is low (high Recall and low Precision), a parameter of *β* = 2 seems insufficient to overcome the poor power of selection. Decreasing this parameter could improve results. However, such too sparse (or too dense) graphs are not informative due to their scarce (or plentiful) number of edges and are not inferred in practice.

#### 3.2.4 Complexity

Even for large-sized networks, BRANE Clust running times remain negligible with respect to weights computation. Networks of size 100 are obtained in few milliseconds while networks composed of 1000 to 5000 nodes are inferred in 1s to 15s. Running times are obtained using an Intel i7-3740QM @ 2.70GHz / 8 Gb RAM and Matlab 2011b. The costlier step is the *random walker* computation. Since the linear system is sparse, implementations with conjugate gradient drastically reduce the complexity, of at most 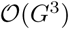, where *G* is the number of nodes.

### 3.3 Inference and insights on predictions

From a biological view point, network inference methods should be evaluated in terms of reliability and predictive power as well. Hence, in addition to numerical performance (AUPR), we analyze the specific additional information acquired by BRANE Clust on real data. For this purpose, we used GENIE3 weights from the third DREAM5 dataset, a compendium of transcriptomic data from *Escherichia coli*. The BRANE Clust network with the best compromise in size and improvement is composed of 236 edges. We compare it to the equally-sized T-GENIE3 network, against the validation.

BRANE Clust infers more true edges than T-GENIE3: 106 compared to 92 (Precision of 45% vs.39%). Amidst 43 edges specific to BRANE Clust, 18 are known to be true links, while only 4 are correctly inferred by T-GENIE3 alone. We are interested in the predictive power of our inference. We thus assess the biological interpretation of the remaining specific 25 novel interactions given by BRANE Clust. We use the STRING database [34]. It contains both known and predicted protein-protein interactions (direct or indirect) from 2031 organisms. Interactions are derived from five main sources: genomic context predictions, high-throughput experiments, (conserved) co-expression, automated textmining and previous knowledge in databases. Several criteria address evidence suggesting a functional link: co-occurrence across genomes (Co-O), co-expression (Co-E), co-mentioned in PubMed abstracts (Co-M), neighborhood in the genome (N), gene fusion, experimental and biochemical data and association in curated databases. A combination of their probabilities, corrected from the chance of randomly observing an interaction, leads to a combined score (CS) per link [35]. Table 3 summarizes significant CS and their associated criteria (Co-O, Co-E, Co-M and N). In addition to quasi-surely well-predicted links, some predicted edges make more sense in terms of grouping targets than in terms of regulatory links. For example, predicted targets for TF *cbl* are *yciW, cysD, cysM, cysA, cysI*. Although links between these genes and the TF *cbl* are not confirmed, genes *cysD, cysM, cysA, cysI* are known to belong to the same operon. In addition, links *yciW-cysD, yciW-cysA* and *yciW-cysI* show a CS of 0.975, 0.897 and 0.946 respectively in the STRING database. Despite a non significant CS for *fis-rpsF* and *allS-gcl*, these two links remain interesting. Indeed, in addition to high CS for *rpsF-rpsI* (0.999) and *allS-allA* (0.808), STRING database also reveals high scores for *fis-rpsI* (0.872) and *allA-gcl* (0.965). This transitivity effect, depicted in Figure 7, suggests a high confidence in the predicted links *fis-rpsF* and *allS-gcl*.

**Table 3.** Significant scores given by the inference-independent STRING database. They reveal plausible functional links from BRANE Clust prediction. Co-O, Co-E, Co-M, N: co-occurrence across genomes, co-expression, co-mentioned in PubMed abstract, neighborhood criteria (resp.). CS: combined score.

| Links | Co-O | Co-E | Co-M | N | CS |
| --- | --- | --- | --- | --- | --- |
| *glcC-mhpR* | - | 0.670 | 0.116 | - | 0.745 |
| *rhaR-rhaT* | 0.699 | 0.678 | 0.867 | - | 0.985 |
| *galS-mglA* | - | 0.915 | 0.403 | 0.370 | 0.975 |
| *ybjG-deoR* | - | 0.149 | - | 0.671 | 0.708 |

**Figure 7.**
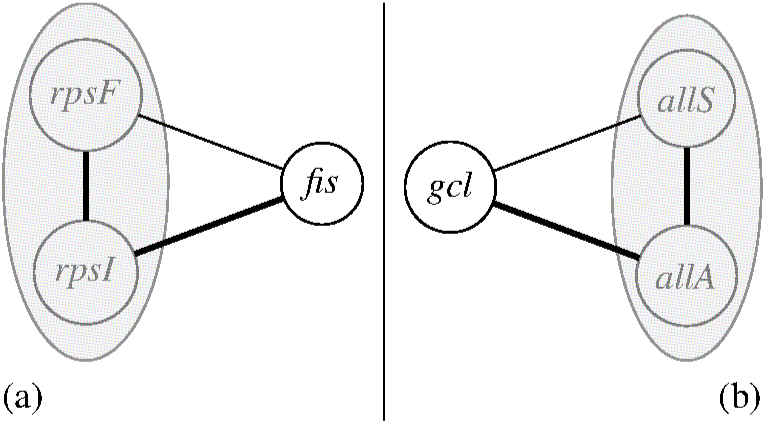
Interpretation for *fis-rpsF* (a) and *allS-gcl* (b) links. Thick edges have significant STRING CS. Normal edges are inferred by BRANE Clust. Genes in operon are embraced in the gray ellipse.

In addition to a better inference of true edges, these results indicate a promising predictive power for BRANE Clust.

### 3.4 Discussions

BRANE Clust results exhibit significant improvements in terms of objective metric (AUPR) and biological interpretation including novel insights in predictions.

On the one hand, in the manner of other methods primarily aimed at inference (e.g. SIMoNe [18]), BRANE Clust improves network construction with gene clustering information. The resulting partitioning being only a secondary product, its evaluation and comparison *per se* is not straightforward. Due to frail inference results for SIMoNe on DREAM4 (with AUPR ten times smaller than those given in Table 1), comparisons — deemed not relevant— are not presented. In addition, graphical models (like SIMoNe) may become extremely costly. Working on gene-to-gene covariance matrices, their 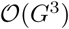-type complexity induces computational difficulties on large-sized networks. As in Section 3.3, it should be judicious to compare the 236-edge network inferred by SIMoNe to the one obtained with BRANE Clust from the third dataset of DREAM5. The resulting clustering of SIMoNe and BRANE Clust may also be analyzed. Unfortunately, with our computational capabilities, SIMoNe failed to return a 236-edge network and the underlying estimated clustering. These limitations did not allow us to pursue the comparison.

On the other hand, WGCNA determines a clustering from a correlation network [25]. Its correlation values are often concentrated in an extremely small interval close to 1. Hence, neither classical nor improved thresholding (e.g. BRANE Clust) is well suited to inference. However, clusters may be more informative — although in a different way — than the network itself. BRANE Clust clustering is compared with WGCNA and X-means clustering [36]. The latter, an extension to K-means [37], [38] with an optimal number of classes, is not specific to biological applications, yet was used recently [39], [40] in this context. Partitions are graded pair-wise, using the Variation of Information (VI) [41], a metric closely related to mutual information. It indicates the closeness of clustering: identical partitioning yields null VI. Our modules (genes arranged around TFs) differ from those in WGCNA or X-means. WGCNA provides 18 modules, X-means 17 clusters, and 322 for BRANE Clust partitioning according to Section 3.3.

Hence, we expect a poor pairwise overlap, confirmed in Figure 8 with significantly non-null VI measures.

**Figure 8.**
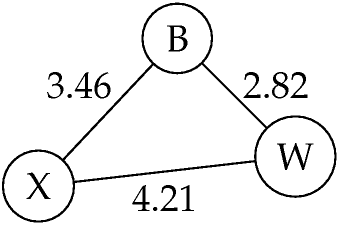
Intrinsic clustering evaluation: pairwise VI measures for BRANE Clust (B), WGCNA (W) and X-means (X).

However, with a closer number of clusters, WGCNA and X-means surprisingly exhibit the largest VI (4.21), thus the least similarity. The best partition overlap (2.82) is observed between WGCNA and BRANE Clust, despite the gap in cluster amount. An external validation with biologically-sound groups of genes from a validated database may be more pertinent. It is built from operons^1^ identified in RegulonDB [42]. This primary database for transcriptional regulation in *Escherichia coli* K-12 contains manually curated knowledge from original scientific publications. All significant operons, containing at least 5 genes, compose the ground truth. It splits a subset of 803 genes into 123 groups. We compare this partitioning to those of BRANE Clust, WGCNA and X-means on the same gene subset in Table 4. A smaller VI (higher similarity) is found for BRANE Clust, suggesting that its partitioning is nigher in terms of operon structure.

**Table 4.** External clustering/operon evaluation: VI measures for BRANE Clust, WGCNA and X-means vs RegulonDB.

|  | BRANE Clust | WGCNA | X-means |
| --- | --- | --- | --- |
| # of clusters | 90 | 18 | 17 |
| VI (vs RegulonDB) | 1.05 | 1.10 | 1.14 |

## 4 CONCLUSIONS

We propose a GRN post-processing tool for classical network thresholding refinement. From a complete weighted network (obtained from any network inference method) BRANE Clust favors edges both having higher weights (as in standard thresholding) and linking nodes belonging to a same cluster. It relies on an optimization procedure. It computes an optimal gene clustering (random walker algorithm) and an optimal edge selection jointly. The introduction of a clustering step in the edge selection process improves gene regulatory network inference. This is demonstrated on both synthetic (five networks of DREAM4 and network 1 of DREAM5) and real (network 3 of DREAM5) data. These conclusions are drawn after comparing classical thresholding on CLR and GENIE3 networks to our proposed post-processing. Significant improvements in terms of Area Under Precision-Recall curve are obtained. The predictive power on real data yields promising results: predicted links specific to BRANE Clust reveal plausible biological interpretation. GRN approaches that produce a complete weighted network to prune could benefit from BRANE Clust post-processing.

In this work, the inference is driven by the clustering, but the graph inference does not influence the clustering. Although the number of clusters is not directly biologically relevant, it may carry useful information in terms of co-expressed groups of genes. The design of novel cluster merging strategies to obtain more relevant gene groupings and networks thus provides a future perspective.

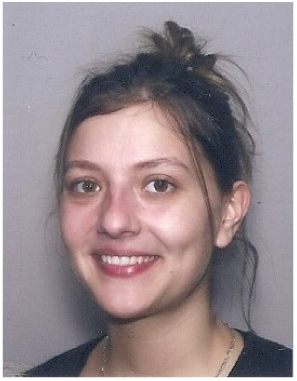

**Aurélie Pirayre** Aurélie Pirayre received the State Engineering degree in bio-sciences from Institut Supérieur des Bio-sciences de Paris (ISBS) in 2013. She is completing a PhD student in signal processing at Université Paris-Est and IFP Energies nouvelles (IFPEN). Her research interests in bioinformatics include gene network inference problems from transcriptomic data. She focuses on aspects of graph theory, penalized criteria, discrete and continuous optimization.

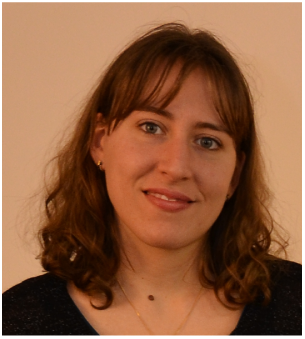

**Camille Couprie** Camille Couprie earned a PhD in computer science at the Université Paris Est and ESIEE Paris, advised by Professors Laurent Najman and Hugues Talbot in 2011, and awarded by the CNRS, DGA and EADS. Then she was postdoctoral researcher at the Courant Institute of Mathematical Sciences at New York University with Professor Yann LeCun, working on semantic segmentation. In 2013 she joined IFP Energies nouvelles in France to work on signal and image processing problems applied to energy problematics. Since 2015 she is a research scientist at Facebook Artificial Intelligence Research in Paris, focusing on the learning of image representations with limited supervision.

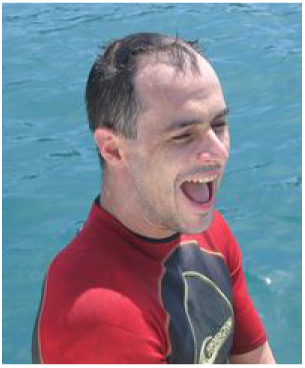

**Laurent Duval** Laurent Duval (S’98–M’00) received the State Engineering degree in electrical engineering from Supélec, Gif-sur-Yvette, France, and the Master (DEA) in pure and applied mathematics from Université de Metz, France, in 1996. He received in 2000 the Ph. D. degree from the Université Paris-Sud (XI), Orsay, France, in the area of seismic data compression. In 1998, he was a research assistant in the MDSP Lab in Boston University, MA, USA (Truong Q. Nguyen’s team). He now works on signal processing and image analysis research in several energy related fields (chemistry, geosciences, biotechnology and transportation) at IFP Energies nouvelles (IFPEN).

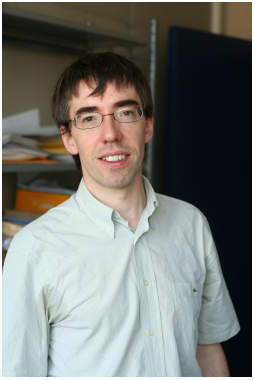

**Jean-Christophe Pesquet** Jean-Christophe Pesquet received the engineering degree from Supelec, Gif-sur-Yvette, France, in 1987, the Ph.D. degree from the University Paris-Sud (XI), Paris, France, in 1990, and the Habilitation à Diriger des Recherches from the University Paris-Sud in 1999. From 1991 to 1999, he was a Maître de Conférences at the University Paris-Sud, and a Research Scientist at the Laboratoire des Signaux et Systemès, Centre National de la Recherche Scientifique (CNRS), Gif-sur-Yvette. From 1999 to 2016, he was a Professor with the University Paris-Est Marne-la-Vallée, France and from 2012 to 2016, he was the Deputy Director of the Laboratoire d’Informatique of the university (UMR-CNRS 8049). He is currently a Professor (exceptional class) with CentraleSupelec, University Paris-Saclay and a Research Scientist at the Center for Visual Computing (INRIA). He is also a senior member of the Institut Universitaire de France and an IEEE Fellow. His research interests include multiscale analysis, statistical signal processing, inverse problems, imaging, and optimization methods with applications to data sciences.

## REFERENCES

[1] S. Klamt, U.-U. Haus, and F. Theis, “Hypergraphs and cellular networks,” PLoS Comput. Biol., vol. 5, no. 5, 2009. [Online]. Available: http://dx.doi.org/10.1371/journal.pcbi.1000385

[2] S. Croset, J. Rupp, and M. Romacker, “Flexible data integration and curation using a graph-based approach,” Bioinformatics, vol. 32, no. 6, pp. 918–925, 2016. [Online]. Available: http://dx.doi.org/10.1093/bioinformatics/btv644

[3] V. Bonnici, F. Busato, G. Micale, N. Bombieri, A. Pulvirenti, and R. Giugno, “APPAGATO: an APproximate PArallel and stochastic GrAph querying TOol for biological networks,” Bioinformatics, vol. 32, no. 14, pp. 2159–2166, 2016. [Online]. Available: http://dx.doi.org/10.1093/bioinformatics/btw223

[4] J. Chen, A. O. Hero, and I. Rajapakse, “Spectral identification of topological domains,” Bioinformatics, vol. 32, no. 14, pp. 2151–2158, 2016. [Online]. Available: http://dx.doi.org/10.1093/bioinformatics/btw221

[5] D. Marbach, R. J. Prill, T. Schaffter, C. Mattiussi, D. Floreano, and G. Stolovitzky, “Revealing strengths and weaknesses of methods for gene network inference,” Proc. Nat. Acad. Sci. U.S.A., vol. 107, no. 14, pp. 6286–6291, Apr. 2010. [Online]. Available: http://dx.doi.org/10.1073/pnas. 0913357107

[6] D. Marbach, J. C. Costello, R. Küffner, N. M. Vega, R. J. Prill, D. M. Camacho, K. R. Allison, The DREAM5 Consortium, M. Kellis, J. J. Collins, and G. Stolovitzky, “Wisdom of crowds for robust gene network inference,” Nat. Meth., vol. 9, no. 8, pp. 796–804, 2012. [Online]. Available: http://www.nature.com/nmeth/journal/vaop/ncurrent/full/nmeth.2016.html

[7] Z. Kurt, N. Aydin, and G. Altay, “A comprehensive comparison of association estimators for gene network inference algorithms,” Bioinformatics, vol. 30, no. 15, pp. 2142–2149, Aug. 2014. [Online]. Available: http://dx.doi.org/10.1093/bioinformatics/btu182

[8] Z.-P. Liu, “Reverse engineering of genome-wide gene regulatory networks from 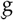ene expression data,” Curr. Genom., vol. 16, no. 1, p. 3âAŞ22, Jan 2015. [Online]. Available: http://dx.doi.org/10.2174/1389202915666141110210634

[9] A. J. Butte and I. S. Kohane, “Mutual information relevance networks: Functional genomic clustering using pairwise entropy measurements,” in Pac. Symp. Biocomput., R. B. Altman, A. K. Dunker, L. Hunter, K. Lauderdale, and T. E. Klein, Eds., vol. 5, Hawaii, HI, USA, Jan. 4-9, 2000, pp. 415–429.

[10] J. J. Faith, B. Hayete, J. T. Thaden, I. Mogno, J. Wierzbowski, G. Cottarel, S. Kasif, J. J. Collins, and T. S. Gardner, “Large-scale mapping and validation of Escherichia coli transcriptional regulation from a compendium of expression profiles,” PLoS Biol., vol. 5, no. 1, pp. 54–66, 2007.

[11] P. E. Meyer, F. Lafitte, and G. Bontempi, “minet: A R/Bioconductor package for inferring large transcriptional networks using mutual information,” Bioinformatics, vol. 9, pp. 461+, 2008.

[12] A. A. Margolin, I. Nemenman, K. Basso, C. Wiggins, G. Stolovitzky, R. Dalla Favera, and A. Califano, “ARACNE: an algorithm for the reconstruction of gene regulatory networks in a mammalian cellular context,” BMC Bioinformatics, vol. 7 (Suppl. 1), no. 5, p. S7, 2006.

[13] X. Zhang, K. Liu, Z.-P. Liu, B. Duval, J.-M. Richer, X.-M. Zhao, J.-K. Hao, and L. Chen, “NARROMI: a noise and redundancy reduction technique improves accuracy of gene regulatory network inference,” Bioinformatics, vol. 29, no. 1, pp. 106–113, 2013. [Online]. Available: http://bioinformatics.oxfordjournals.org/content/29/1/106.abstract

[14] H. Toh and K. Horimoto, “Inference of a genetic network by a combined approach of cluster analysis and graphical Gaussian modeling,” Bioinformatics, vol. 18, no. 2, pp. 287–297, Feb. 2002. [Online]. Available: http://dx.doi.org/10.1093/bioinformatics/18.2.287

[15] N. Friedman, “Inferring cellular networks using probabilistic graphical models,” Science, vol. 303, no. 5659, pp. 799–805, Feb. 2004. [Online]. Available: http://dx.doi.org/10.1126/science.1094068

[16] H. Lähdesmäki, S. Hautaniemi, I. Shmulevich, and O. Yli-Harja, “Relationships between probabilistic Boolean networks and dynamic Bayesian networks as models of gene regulatory networks,” Signal Process., vol. 86, no. 4, pp. 814–834, Apr. 2006. [Online]. Available: http://dx.doi.org/10.1016/j.sigpro.2005.06.008

[17] P. Li, C. Zhang, E. J. Perkins, P. Gong, and Y. Deng, “Comparison of probabilistic Boolean network and dynamic Bayesian network approaches for inferring gene regulatory networks,” BMC Bioinformatics, vol. 8 (Suppl. 7), p. S13, 2007. [Online]. Available: http://dx.doi.org/10.1186/1471-2105-8-S7-S13

[18] J. Chiquet, A. Smith, G. Grasseau, C. Matias, and C. Ambroise, “SIMoNe: Statistical Inference for MOdular NEtworks,” Bioinformatics, vol. 25, no. 3, pp. 417–418, 2009.

[19] A. Wiesel, Y. C. Eldar, and A. O. Hero III, “Covariance estimation in decomposable Gaussian graphical models,” IEEE Trans. Signal Process., vol. 58, no. 3, pp. 1482–1492, Mar. 2010. [Online]. Available: http://dx.doi.org/10.1109/TSP.2009.2037350

[20] Z.-P. Liu, H. Wu, J. Zhu, and H. Miao, “Systematic identification of transcriptional and post-transcriptional regulations in human respiratory epithelial cells during influenza a virus infection,” BMC Bioinformatics, vol. 15, no. 1, p. 336, 2014. [Online]. Available: http://dx.doi.org/10.1186/1471-2105-15-336

[21] Z.-P. Liu, C. Wu, H. Miao, and H. Wu, “RegNetwork: an integrated database of transcriptional and post-transcriptional regulatory networks in human and mouse,” Database, vol. 2015, p. bav095, 2015. [Online]. Available: http://dx.doi.org/10.1093/database/bav095

[22] V. A. Huynh-Thu, A. Irrthum, L. Wehenkel, and P. Geurts, “Inferring regulatory networks from expression data using tree-based methods,” PLoS One, vol. 5, no. 9, p. e12776, Sep. 2010. [Online]. Available: http://dx.doi.org/10.1371/journal.pone.0012776

[23] A. Pirayre, C. Couprie, F. Bidard, L. Duval, and J.-C. Pesquet, “BRANE Cut: biologically-related a priori network enhancement with graph cuts for gene regulatory network inference,” BMC Bioinformatics, vol. 16, no. 1, p. 369, Dec. 2015. [Online]. Available: http://dx.doi.org/10.1186/s12859-015-0754-2

[24] M. Banf and S. Y. Rhee, “Enhancing gene regulatory network inference through data integration with markov random fields,” Sci. Rep., 7:41174, Feb. 2017. [Online]. Available: http://dx.doi.org/10.1038/srep41174

[25] P. Langfelder and S. Horvath, “WGCNA: an R package for weighted correlation network analysis,” BMC Bioinformatics, vol. 9, no. 1, p. 559, 2008. [Online]. Available: http://dx.doi.org/10.1186/1471-2105-9-559

[26] M. E. J. Newman, “Communities, modules and large-scale structure in networks,” Nat. Phys., vol. 8, no. 1, pp. 25–31, 2012. [Online]. Available: http://dx.doi.org/10.1038/nphys2162

[27] E. Segal, M. Shapira, A. Regev, D. Pe’er, D. Botstein, D. Koller, and N. Friedman, “Module networks: identifying regulatory modules and their condition-specific regulators from gene expression data,” Nat. Genet., vol. 34, no. 2, pp. 166–176, May 2003. [Online]. Available: http://dx.doi.org/10.1038/ng1165

[28] A. Joshi, R. De Smet, K. Marchal, Y. Van de Peer, and T. Michoel, “Module networks revisited: computational assessment and prioritization of model predictions.” Bioinformatics, vol. 25, no. 4, pp. 490–496, 2009. [Online]. Available: http://dx.doi.org/10.1093/bioinformatics/btn658

[29] S. Roy, S. Lagree, Z. Hou, J. A. Thomson, R. Stewart, and A. P. Gasch, “Integrated module and gene-specific regulatory inference implicates upstream signaling networks,” PLoS Comput. Biol., vol. 9, no. 10, p. e1003252, Oct. 2013. [Online]. Available: http://dx.doi.org/10.1371/journal.pcbi.1003252

[30] A. Pirayre, C. Couprie, L. Duval, and J.-C. Pesquet, “Graph inference enhancement with clustering: Application to gene regulatory network reconstruction,” in Proc. Eur. Sig. Image Proc. Conf., Nice, France, Aug. 31-Sep. 4, 2015, pp. 2406–2410. [Online]. Available: http://dx.doi.org/10.1109/EUSIPCO.2015.7362816

[31] L. Grady, “Random walks for image segmentation,” IEEE Trans. Pattern Anal. Mach. Intell., vol. 28, no. 11, pp. 1768–1783, Nov.2006. [Online]. Available: http://dx.doi.org/10.1109/TPAMI.2006.233

[32] D. Marbach, T. Schaffter, C. Mattiussi, and D. Floreano, “Generating realistic in silico gene networks for performance assessment of reverse engineering methods,” J. Comput. Biol., vol. 16, no. 2, pp. 229–239, Feb. 2009. [Online]. Available: http://dx.doi.org/10.1089/cmb.2008.09TT

[33] T. Schaffter, D. Marbach, and D. Floreano, “GeneNetWeaver: in silico benchmark generation and performance profiling of network inference methods,” Bioinformatics, vol. 27, no. 16, pp. 2263–2270, 2011. [Online]. Available: http://dx.doi.org/10.1093/bioinformatics/btr373

[34] A. Franceschini, D. Szklarczyk, S. Frankild, M. Kuhn, M. Simonovic, A. Roth, J. Lin, P. Minguez, P. Bork, C. von Mering, and L. J. Jensen, “STRING v9.1: protein-protein interaction networks, with increased coverage and integration,” Nucleic Acids Res., vol. 41, pp. D808–D815, 2013.

[35] C. von Mering, L. J. Jensen, B. Snel, S. D. Hooper, M. Krupp, M. Foglierini, N. Jouffre, M. A. Huynen, and P. Bork, “STRING: known and predicted protein-protein associations, integrated and transferred across organisms,” Nucleic Acids Res., vol. 33 (Suppl. 1), pp. D433–D437, Jan. 2005. [Online]. Available: http://dx.doi.org/10.1093/nar/gki005

[36] D. Pelleg and A. Moore, “X-means: Extending K-means with efficient estimation of the number of clusters,” in Proc. Int. Conf. Mach. Learn., Stanford, CA, USA, Jun. 29-Jul. 2 2000, pp. 727–734.

[37] H. Steinhaus, “Sur la division des corps matériels en parties,” Bull. Acad. Polon. Sci., vol. Cl. III — Vol. IV, no. 12, pp. 801–804, 1956.

[38] J. MacQueen, “Some methods for classification and analysis of multivariate observations,” in Proc. Fifth Berkeley Symp. Math. Statist. Prob., 1967, pp. 281–297.

[39] M. Wang, N. Jiang, T. Jia, L. Leach, J. Cockram, R. Waugh, L. Ramsay, B. Thomas, and Z. Luo, “Genome-wide association mapping of agronomic and morphologic traits in highly structured populations of barley cultivars,” Theor. Appl. Genet., vol. 124, no. 2, pp. 233–246, Sep. 2012. [Online]. Available: http://dx.doi.org/10.1007/s00122-011-1697-2

[40] A. Halleran, S. Clamons, and M. Saha, “Transcriptomic characterization of an infection of Mycobacterium smegmatis by the cluster A4 mycobacteriophage Kampy,” PLoS One, vol. 10, no. 10, p. e0141100, Oct. 2015. [Online]. Available: http://dx.doi.org/10.1371/journal.pone.0141100

[41] M. Meilă, “Comparing clusterings—an information based distance,” J. Multivariate Anal., vol. 98, no. 5, pp. 873–895, May 2007. [Online]. Available: http://dx.doi.org/10.1016/j.jmva.2006.11.013

[42] S. Gama-Castro, H. Salgado, A. Santos-Zavaleta, D. Ledezma-Tejeida, L. Muñiz-Rascado, J. S. García-Sotelo, K. Alquicira-Hernández, I. Martínez-Flores, L. Pannier, J. A. Castro-Mondragón, A. Medina-Rivera, H. Solano-Lira, C. Bonavides-Martínez, E. Pérez-Rueda, S. Alquicira-Hernández, L. Porrón-Sotelo, A. López-Fuentes, A. Hernández-Koutoucheva, V. Del Moral-Chávez, F. Rinaldi, and J. Collado-Vides, “RegulonDB version 9.0: high-level integration of gene regulation, coexpression, motif clustering and beyond,” Nucleic Acids Res., vol. 44, no. D1, pp. D133–D143, 2016. [Online]. Available: http://dx.doi.org/10.1093/nar/gkv1156

